# The influence of drug dosing regimen on extinction and adaptation during experimental evolution in *Candida albicans*

**DOI:** 10.64898/2026.10.29.762568

**Authors:** Parul Sethi, Erica Pouteau, Aruni Sumanarathne, Aleeza C. Gerstein

## Abstract

The antifungal azole drug fluconazole (FLC) is frequently prescribed for treating *Candida albicans* infections. Azole drugs are typically fungistatic—they inhibit cell growth rather than kill susceptible cells within a population. Therefore, fungal populations can adapt to fluconazole through increased drug resistance or increased drug tolerance (the ability of a subpopulation of susceptible cells to grow slowly at inhibitory drug concentrations). To our knowledge, no prior experiments have examined how different factors influence extinction propensity, although previous *in vitro* studies show that both drug concentration and strain background can influence adaptation. To determine how the drug dosing regimen, namely fluconazole concentration and transfer duration (i.e., the amount of time between transfer into fresh medium), influenced extinction and adaptation in *Candida albicans*, we conducted a fully factorial series of short *in vitro* evolution experiments. Dosing regimen significantly influenced extinction: all replicates evolved at all drug concentrations survived the 48 h and 72 h transfer experiments, yet the extinction rate increased sharply from low to high drug concentration with 24 h transfers. Increases in drug tolerance were frequent regardless of the dosing regimen. In contrast, drug resistance and growth rate both tended to decrease following evolution at higher drug concentrations. Fluconazole-evolved replicates exhibited frequent increases in cross-tolerance and decreases in cross-resistance in a panel of other azole drugs. Genome size changes were common in some strain backgrounds evolved to higher fluconazole concentrations, regardless of transfer times. Combined, our results indicate that many experimental factors can influence the trajectory of adaptation, and show that extinction in a fungistatic drug occurs readily at high drug concentration with rapid transfers.

## INTRODUCTION

*Candida albicans* is a global opportunistic fungal pathogen and a common member of the human microbiome across diverse body sites. It was recently named to the WHO Fungal Priority Pathogen List as one of the top priority fungal pathogens (World Health Organization. Antimicrobial Resistance Division et al., 2022). As fungal eukaryotic cells are very similar to human cells, developing antifungal drugs is challenging. *C. albicans* infections are commonly limited to three major classes of antifungal drugs, the polyenes, azoles, and echinocandins (Perfect, 2017), which differ in their cellular targets and mode of action. The most common are the azole drugs, which include two subclasses, the triazoles and the imidazoles. Triazoles such as fluconazole (FLC), posaconazole (POS), and itraconazole (ITZ) contain three nitrogen atoms in their cyclic ring, while the imidazole group, including clotrimazole (CTR) and miconazole (MCZ) contain two nitrogen atoms in their cyclic rings. FLC is often the preferred treatment for *Candida* infections because of its bioavailability, low cost, and low toxicity (de Oliveira Santos et al., 2018). Daily FLC doses ranging between 150-800 mg are administered to treat different types of candidiasis, yielding serum concentrations around 7.5–60.5 μg/mL (Govindarajan et al., 2023). Thus, physiological concentrations of FLC vary widely across different tissues and body fluids, and this variation may influence the evolutionary dynamics of the focal fungal populations.

FLC acts by binding to and inhibiting the activity of the enzyme lanosterol 14-demethylase, which is involved in the biosynthetic pathway of the principal sterol component of the fungal cell membrane, ergosterol. This binding disrupts the cell membrane and leads to the accumulation of toxic C14α-methyl sterols and elevated levels of reactive oxygen species (ROS) (Chaud et al., 2020; Pfaller et al., 2013). Many different FLC resistance mechanisms have been documented among clinical isolates and in the lab, with the majority of mutations involving alterations in the azole drug target 14-α-demethylase (*ERG11*, also referred to as *CYP51* in other species), resulting in overexpression of the drug efflux pumps (ABC transporters, *CDR1* and *CDR2*, and *MDR1*), and in the transcription factors that regulate the ergosterol biosynthesis pathway (*UPC2*, *TAC1* and *MRR1*) (Bédard et al., 2025). In addition, chromosomal copy number changes that include the genomic location of many of these genes, such as chromosomes III, IV, V, and R, have also been documented (Bédard et al., 2025).

Fluconazole is typically fungistatic, that is, it inhibits cell growth rather than killing susceptible cells within the population. In addition to acquisition of drug resistance (i.e., an increase in the minimum inhibitory concentration at which the majority of cells in the population are able to grow well at), changes in drug tolerance (an increased size in the subpopulation of susceptible cells that are able to grow slowly at drug concentrations above the MIC) have also been observed (Berman & Krysan, 2020; Rosenberg et al., 2018; Yang & Berman, 2024). The mechanism of tolerance is distinct from resistance (Gerstein et al., 2016; Rosenberg et al., 2018; Salama & Gerstein, 2022) and seems to be polygenic (Berman & Krysan, 2020; Schutz et al., 2025). Existing studies suggest that tolerance is associated with multiple signal transduction pathways (Rosenberg et al., 2018), including Hsp90, calcineurin, and the protein kinase C cascade (Chen et al., 2023), secretory trafficking (Gervais, manuscript in prep), ergosterol biosynthesis and protein homeostasis (Berman & Krysan, 2020; Yau et al., 2023). Some evidence suggests that persistent candidemia may result from increased FLC tolerance that hinders successful treatment, rather than increased resistance (Levinson et al., 2021; Rosenberg et al., 2018). Many more clinical studies are needed to determine the interplay between drug resistance and tolerance in causing treatment failure.

Experimental evolution with microbial species that grow rapidly *in vitro* has become a widely used method for testing hypotheses about the evolutionary process (Kassen, 2024; Van den Bergh Bram et al., 2018). Due to their short generation time, relatively small genomes, and the ability to preserve a fossil record, it is possible to easily propagate many parallel replicate populations over hundreds or thousands of generations on short time scales under well-controlled conditions. Evolved replicates can then be compared against their progenitor (and against each other) using high-throughput phenotypic and genotypic methods to pinpoint the traits and genetic changes that underpin adaptation. The most common method is batch culture evolution starting from a single clone, which involves repeatedly diluting the evolving populations at regular intervals into fresh medium to supply nutrients and remove toxic end products. Many different variables that could influence evolutionary dynamics can be controlled by the researcher (Izutsu et al., 2024; Lenski, 2023): the species and specific isolate(s) to use as the ancestor, the specific medium (including, e.g., different levels of carbon or nutrients, pH, temperature, addition of stress and the degree of stress), temperature, culture vessel (which influences the volume of medium and thus maximal population size as well as the degree of aeration), the dilution factor (the proportion of culture/cells that are transferred), the time between transfers, and the number of total generations. Taken alone or in combination, these factors influence the rate at which new mutations enter the population and the efficacy of selection, which in turn affects the pace of fitness change and the rate of adaptation and genotypic and phenotypic parallelism. Experimental evolution, particularly with an “evolve-and-resequence” approach (Cooper, 2018) can thus be used to study the fundamental properties that dictate whether and how populations evolve to the environment they find themselves in.

The first *in vitro* experiment exposed *C. albicans* replicates to stepwise increases in fluconazole concentration and found that MIC increased in all replicates, though the rate of change and the terminal MIC varied. Since then, many additional experiments to characterize the acquisition of drug resistance have been conducted. These experiments have examined evolution of drug resistance to a single level of fluconazole, typically at or below the MIC, with transfers done after a set number of hours (typically every 48 or 72 h) or were designed to study the evolution of drug resistance to a stepwise increase in fluconazole levels from low to high (with transfer done either at set times or once a specific optical density has been reached; (Bing et al., 2020; Burrack et al., 2022; Cowen et al., 2000; Handelman & Osherov, 2022; Mount et al., 2018; Rybak et al., 2017). A handful of studies in recent years have also examined drug tolerance using an experimental evolution framework, revealing that at low drug concentrations, populations typically increase in resistance only up to the drug concentration they were evolved in, while evolution at high drug concentrations selected for increased tolerance (Gerstein & Berman, 2020; Todd et al., 2023; Yang et al., 2023). In addition to fluconazole, increased tolerance was also found when *C. albicans* strains were evolved to high concentrations of miconazole (MCZ) (Guo et al., 2024) or posaconazole (POS) (Kukarudz 2022). Interestingly, some *C. glabrata* isolates evolved from supra-MIC fluconazole levels also increased in tolerance, though many more such isolates became resistant (Zheng et al., 2024). Recent studies have also highlighted the importance of strain background, independent of ancestral MIC, on evolutionary trajectory (Burrack et al., 2022; Gerstein & Berman, 2020; Kukurudz et al., 2022; Todd et al., 2023). Together, these evolution experiments show that multiple drug response phenotypes can change during adaptation to fluconazole and that caution is warranted before extrapolating from a single experiment, as many factors influence evolutionary dynamics.

The majority of *in vitro* evolution experiments have been designed to promote adaptation; indeed, “adaptive laboratory evolution” is often used as a synonym for “experimental evolution” in the microbial literature. As such, in experiments where the stress level is constant, it is typically chosen to be moderate (i.e., sub-inhibitory in the context of antimicrobials). In experiments where the level of stress is increased over time, the pace of stress increase is typically chosen to be slower than the rate at which beneficial mutations enter the population, to provide replicates with the opportunity to adapt (Gerstein & Sethi, 2022). Yet in these experiments, as in nature, extinction of replicates is also a possible outcome. Many experiments have been conducted to identify the conditions necessary for evolutionary rescue, i.e., the process where evolution is required for a population to avoid extinction (Alexander et al., 2014; Gomulkiewicz & Holt, 1995). Generally, extinction occurs if the beneficial mutation rate is lower than the effective population size or if the population size is decreasing at the time of transfer (Van den Bergh et al., 2018), matching the conditions experimenters choose when setting up experimental evolution studies. Extinction of some replicates does, however, occur in some evolution experiments, yet is not always thoroughly reported (Gerstein & Sethi, 2022).

We previously conducted two experimental evolution studies in different drugs, where one of the goals was to examine extinction rates in *C. albicans*. In the posaconazole experiment, we picked a consistent level of drug stress that was above the MIC of all eight ancestral strain backgrounds and an order of magnitude higher than the epidemiological cutoff value. None of the 96 replicates survived when transfers were conducted every 24 h, whereas approximately half survived through 72 h transfers (Kukurudz et al., 2022). In boric acid, we started with a level of drug that was at or below the MIC_50_ of the eight ancestral strains, and increased the level of drug over time. Replicates started to go extinct at 3× the MIC_50,_ and all replicates were extinct by 10× MIC_50_ (Syvolos et al., 2024). In both cases, the rate of extinction was influenced by strain background, independent of ancestral MIC_50_.

Here, we set out to explore the interplay among genetic background, drug concentration, and transfer duration on extinction probability, as well as changes in drug tolerance and resistance. We used a fully factorial design to conduct nine short-term evolution experiments in three levels of drug (0.25 μg/mL FLC, a subinhibitory level of drug; 1 μg/mL FLC, a level of drug near the MIC_50_, and 16 μg/mL FLC, a supraMIC level of drug) and three transfer times (24 h, 48 h, 72 h). For each experiment, we evolved 24 replicates from four *C. albicans* strains belonging to different genetic backgrounds (GC75, SC5314, P78048, P75016). We refer to FLC concentration and transfer time collectively as the “drug dosing regimen”. Our results highlight that transfer duration, particularly at high levels of drug, is critically important for extinction. We also add to the growing body of work showing that drug tolerance and resistance evolve independently of each other, with increases in drug tolerance observed much more frequently than drug resistance across all dosing regimens. Importantly, we also demonstrate that growth rate is not an appropriate proxy for fitness in this context, as growth rate surprisingly decreased over evolutionary time.

## METHODS

### Strains and isolation of ancestral replicates

Four strains of *Candida albicans* belonging to two genotypically distinct clades (clade one and clade four) were used in our study. This included strains with the same parental MIC_50_ for fluconazole (FLC) - GC75 (MIC_50_= 0.0125 µg/mL) (Wu et al., 2007), SC5314 (MIC_50_= 0.0125 µg/mL) (Lockhart et al., 2002), P78048 (MIC_50_= 0.5 µg/mL) (Wu et al., 2007), and P75016 (MIC_50_= 0.5 µg/mL) (Wu et al., 2007). Frozen glycerol stock of each strain was streaked onto agar plates of the standard rich lab medium for yeast, YPD, supplemented with adenine sulfate and uridine to improve growth (2% w/v peptone, 2% w/v yeast extract, 0.00016% w/v adenine sulfate, 0.00008% w/v uridine, 1.8% w/v agar, 1% w/v glucose, 0.1% v/v of each chloramphenicol and ampicillin) and incubated for 48 h at 30 °C. For each strain, three plates were spotted randomly eight times. The 24 colonies closest to each dot were randomly chosen and inoculated into 1 mL of YPD in a deep (2 mL) 96-well culture box (2 mL), followed by incubation at 30 °C, shaking at 350 rpm overnight. From each overnight culture, 50 μL was frozen for each replicate in triplicate in 25% glycerol and stored at −70 °C. These are referred to as the ancestral replicates.

### Experimental evolution experiments

The ancestral replicates were inoculated from frozen and grown in YPD overnight. Each replicate was then diluted 1:1000 in two steps in 1 mL volumes (1:100 then 1:10) into low drug (YPD + 0.25 μg/mL fluconazole, below the MIC_50_), medium drug (YPD + 1 μg/mL fluconazole, at MIC_50_), and high drug (YPD + 16 μg/mL fluconazole, above MIC_50_). 200 uL of the final culture was placed into a round-bottom microtiter plate (Grenier, Ref 650161); to try and minimize cross-contamination among strains, rows A and C were always used for replicates from one strain, and rows F and H were used for a second strain. Sterile distilled water was placed in the remaining rows. Breathe-easier membranes (Electron Microscopy Sciences, PA, United States) were used to seal the microtiter plates, and they were incubated statically at 30 °C for either 24 h, 48 h, or 72 h, depending on the appropriate transfer duration.

After incubation (for 24 h, 48 h, or 72 h), the cultures were mixed by gently pipetting 3-5 times, followed by the same two-step 1:1000 transfer into fresh YPD + FLC. This was done four times for each dosing regimen (FLC concentration × transfer time), i.e., ∼ 40 generations of evolution. After the fourth transfer, 50 μL of culture from each evolved replicate was frozen in quadruplicates in 25% glycerol and stored at −70 °C.

One evolution experiment was thus performed for each of the nine dosing regimens. The experiment nomenclature is FLC concentration-transfer time, e.g., “FLC0.25-24hT” for the experiment conducted in YPD + 0.25 μg/mL FLC with transfers every 24 h. All pipette steps were done with multichannel pipettes unless otherwise stated.

### Extinction

We scored extinction based on our inability to revive evolved replicates after multiple attempts from the freezer stock (this included thawing the tubes completely, centrifugation, and pipetting the liquid from the bottom into fresh YPD). To increase the statistical power of the 24 h extinction data (the only transfer time at which extinction was observed in the initial experiment), we repeated the 24 h transfer experiments at the three drug levels.

### Fluconazole drug response quantification

Antifungal susceptibility and tolerance testing were performed using a disk diffusion assay from 12 replicates evolved at each transfer duration and drug concentration from each strain background (haphazardly chosen). The standardized CLSI guidelines were adapted to more closely mimic the conditions of the experiment. Briefly, 5 μL of ancestral and evolved replicates were inoculated from frozen into 500 μL of YPD in a deep 96-well culture box, covered with a breathe-easier sealing membrane, and incubated at 30 °C shaking at 250-350 rpm for 48 h. The culture in each well was mixed well by pipetting 3-5 times, 200 μL of culture was transferred into a 96-well microtiter plate, and the OD_600_ was measured. Culture from all replicates were standardized to OD_600_ 0.01 in PBS in a deep 96-well culture box. In duplicate, 100 µL of standardized culture was added to a YPD plate and spread with sterile 5 mm glass beads. The beads were removed, and the plates were turned upside down to dry for ∼20 minutes. An antifungal disc was placed in the center of each plate, followed by incubation for 48 h at 30 °C. Ancestral and evolved replicates from the same drug dosing regimen were always conducted on the same experimental day.

Fluconazole disks (25 µg/mL) were purchased (Oxoid Ltd, UK), disks for other drugs were prepared from liquid stock in DMSO added to blank disks (Fisher Scientific) to be consistent with the CLSI guidelines. Clotrimazole disks were prepared by adding 1 mg/mL CTR to a final concentration of 10 µg. Miconazole disks were prepared by adding 2 mg/mL of MCZ to a final concentration of 10 µg. Itraconazole disks were prepared by adding 5 mg/mL ITZ stock to a final concentration of 50 µg. Posaconazole disks were prepared by adding 0.5 mg/mL POS stock to a final concentration of 5 µg.

Photographs of the 48 h incubated plates were taken on a lightbox. The raw images were converted to 8-bit, inverted, and brightness and contrast were adjusted using consistent parameters in ImageJ to maximize the bright white colonies against a dark background. The processed images were analyzed with the *diskImageR* R package, which computationally measures the drug susceptibility as the radius where an 80 % reduction in growth occurs relative to the perimeter of the plate (RAD_80_) and drug tolerance as the fraction of growth above RAD_80_ (FOG_80_) (Gerstein et al., 2016). The two technical replicates were averaged together.

### Quantification of cross-resistance and cross-tolerance

Drug Susceptibility was measured by disk diffusion assays for two technical replicates for 12 biological replicates for each ancestral and evolved strain at FLC0.25 and FLC16. Four susceptibility experiments were performed at four different times. Other drugs tested included POS, ITZ, CTR, and MCZ. Fluconazole (25 µg) disks were purchased (Fisher Scientific, Ottawa ON, Canada). All other disks were prepared according to Clinical and Laboratory Standards Institute (CLSI) guidelines. Clotrimazole disks were prepared by using 1 mg/mL CTR stock in DMSO added to blank susceptibility disks (Fisher Scientific, Ottawa ON, Canada) for a final concentration of 10 µg. Miconazole disks were prepared by using 2 mg/mL MCZ stock in DMSO added to blank susceptibility disks (Fisher Scientific, Ottawa ON, Canada) for a final concentration of 10 µg. Itraconazole disks were prepared by using 5 mg/mL ITZ stock in DMSO added to blank susceptibility disks (Fisher Scientific, Ottawa ON, Canada) for a final concentration of 50 µg. Posaconazole disks were prepared by using 0.5 mg/mL POS stock in DMSO added to blank susceptibility disks (Fisher Scientific, Ottawa ON, Canada) for a final concentration of 5 µg. The protocol for all disk assays was otherwise the same as what was done for fluconazole.

### Growth rate measurement

Growth rates were measured for ancestral and evolved replicates in the evolutionary drug environment in which the replicates were evolved. Briefly, 5 µL of the frozen ancestral and evolved strain replicates were inoculated into 500 µL of liquid YPD in a deep 96-well box, covered with a breathe-easy sealing membrane (Electron Microscopy Sciences, PA, United States), and incubated with shaking (250 – 350 rpm) at 30 °C for 48 h. The cultures were then standardized to OD_600_ 0.02 in YPD, and 100 µL of the standardized culture was added to 100 µL of YPD + double the evolutionary concentration of FLC in round microtiter plates. The plates were covered with a clear breathe-easy sealing membrane and incubated at 30 °C in an Epoch microplate spectrophotometer with continuous shaking. OD_600_ was measured every 15 minutes for 24 h, 48 h, or 72 h, depending on the transfer time of the evolution experiment from which the strain replicates were being analyzed. A custom R script by Dr. Richard Fitzjohn was used to determine the spline with the highest slope from each growth curve to calculate the maximal growth rate per well (Syvolos et al., 2024). The average growth rate between two biological replicates of each strain for each evolution experiment was used for analysis and visualization.

### Genome size estimation

Flow cytometry was used to evaluate whether the genome size of evolved replicates had increased relative to their diploid ancestors. Briefly, 5 µL of ancestral and evolved replicates were inoculated from frozen into 500 µL of liquid YPD in a deep 96-well box, covered with a breathe-easy sealing membrane, and incubated with shaking at 30 °C for 48 h. After 48 h, 10 µL was subcultured into 500 µL of fresh liquid YPD, covered with a breath-easier membrane, and incubated shaking at 30 °C for 4 hours. 200 µL of the subculture was transferred to round bottom 96-well microtiter plates and centrifuged at 1000 x g for 5 minutes (these settings were used for all centrifugation steps). The supernatant was discarded, followed by resuspension of the pellets in 20 µL of 50:50 Tris-EDTA (TE buffer). 180 µL of cold 95% ethanol was slowly added to fix the cells in the resuspended pellet, and the plates were wrapped in aluminum foil and stored at −20 °C. Fixed cells were typically stored for two to three weeks. The fixed culture was thawed, then centrifuged. The supernatant was discarded, and the pellets were washed in 200 µL of TE. The plates were pelleted again and resuspended in 50 µL of 1 mg/mL RNase A solution, followed by overnight incubation at 37 °C. The samples were then centrifuged again, the supernatant was discarded, and 50 µL of TE buffer was added to resuspend the pellets. To this, 50 µL of 1:100 sytox: 50/50 TE solution was added, and the plates were incubated in the dark overnight. The overnight plates were centrifuged, and 200 µL TE was added to the pellets. After mixing, the resuspension was added to flow tubes containing 500 µL TE. Flow cytometry was performed on an SH008S cell sorter (Sony Biotechnology Inc, San Jose, California, United States). 10,000 events were recorded for each population.

The resulting data was analyzed using the *PloidyPeaks* R package (https://github.com/margothenry/PloidyPeaks), which determines the G1 mean of each population. In addition, *PloidyPeaks* assesses whether there is statistical support for a mixed-ploidy population and estimates the number of cells in each, effectively identifying the dominant genome size in each evolved population. The genome size of all ancestral (t0) replicates was always measured at the same time as t4 replicates. The dominant evolved genome size from all populations was divided by the median ancestral genome size and multiplied by two (to set the base ploidy at diploid).

## RESULTS

### Extinction increases with evolutionary drug concentration with 24 h transfers

Replicates from four genetic backgrounds of *C. albicans* were passaged with 1:1000 dilutions every 24 h, 48 h, and 72 h for a total of four transfers in low (YPD + FLC0.25), medium (FLC1), and high (FLC16) drug concentrations. All strain replicates survived to the end of the 48 h and 72 h transfer evolution experiments. In the 24 h transfer experiments, some replicates went extinct across all drug concentrations (Figure 1), with the number surviving decreasing with increasing drug concentration and varying by strain background. GC75 replicates had the highest extinction at all drug levels, while SC5314 replicates had the lowest (Figure 1; fixed-effect model, with the proportion of replicates that survived as the response variable and drug concentration and strain as categorical variables, p-value obtained through two-way ANOVA; drug concentration: F_2,6_ = 25.63, p = 0.001; strain: F_3, 6_ = 5.02, p = 0.045 for strain).

**Figure 1.**
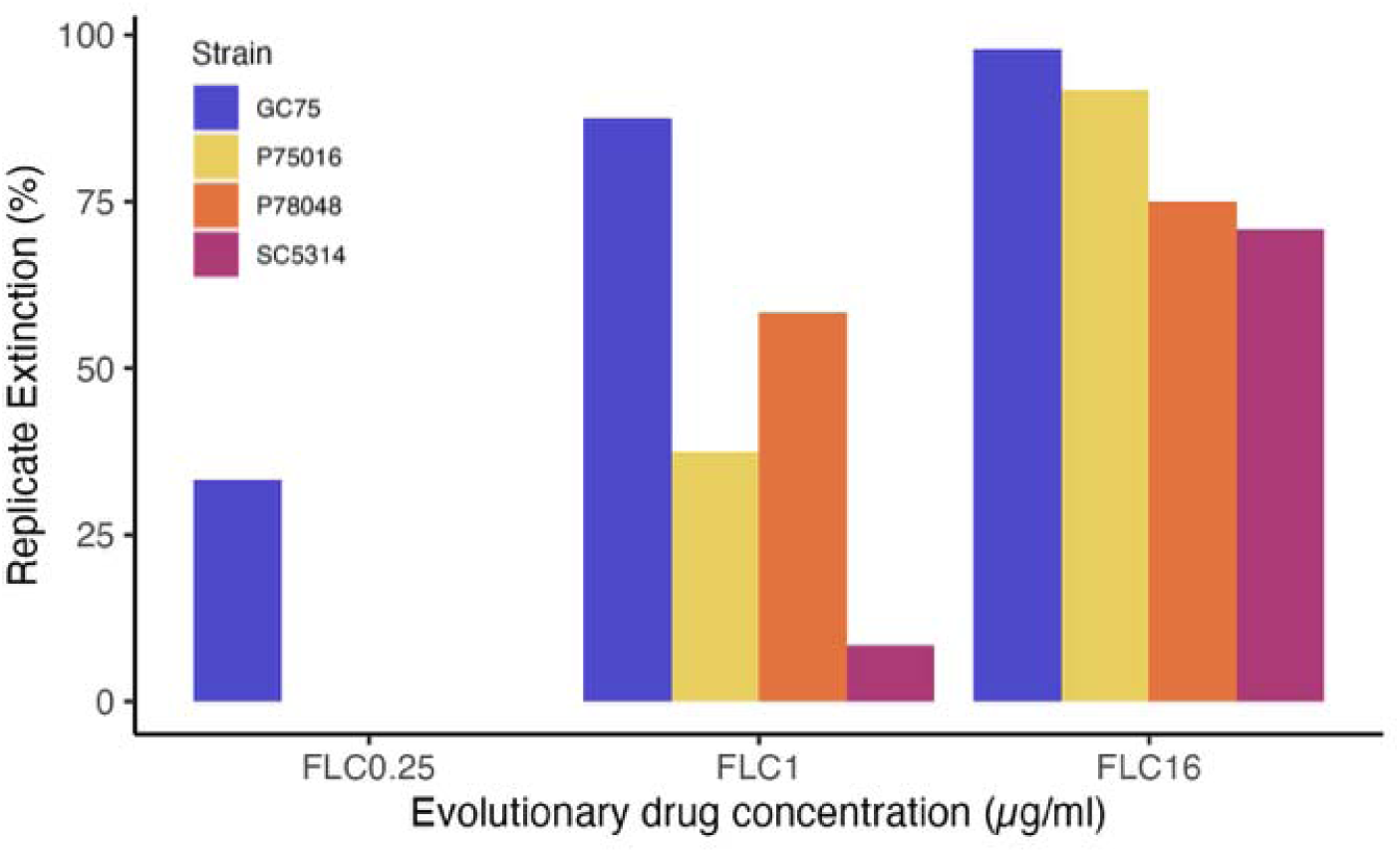
Extinction during 24 transfers increases the drug concentration increased. Replicates evolved to low (FLC0.25), medium (FLCl), and high (FLC16) fluconazole concentrations was calculated from 24 evolved replicates from each strain background evolved in FLC0.25 and FLCl and 48 evolved replicates from each strain background in FLC16.

### Drug tolerance increased while drug resistance and growth rate decreased across the drug dosing regimens

The overall phenotypic change in evolved replicates was remarkably consistent across dosing regimens and strain backgrounds; drug tolerance increased while drug resistance and growth rate decreased (Figure 2). Considering the mean difference among evolved replicates within a strain compared to the ancestral replicates, drug tolerance and drug resistance were significantly influenced by both evolutionary drug concentration and transfer time (but not their interaction), while growth rate was only significantly influenced by drug concentration. Statistical significance was investigated through linear mixed-effects models implemented using the lmer R package (Kuznetsova et al., 2017), with the p-values evaluated through a Type II ANOVA with Satterthwaite’s method; the mean difference for each trait was the response variable, with drug concentration, transfer time, and their interaction as the predictor variables and strain as a random effect: drug tolerance---drug concentration, F_2, 26_ = 15.03, p < 0.0001; transfer time, F_2,26_ = 4.98, = 0.015; drug × transfer time, F_4,26_ =2.42, p = 0.074; drug resistance---drug concentration, F_2, 26_ = 6.66, p = 0.0046; transfer time, F_2,26_ = 4.28, = 0.025; drug × transfer time, F_4, 26_ =0.62, p = 0.65; growth rate---drug concentration, F_2, 23.3_ = 2.11, p = 0.1437; transfer time, F_2,23.3_ = 23.48, p < 0.0001; drug × transfer time, F_4, 23.3_ = 1.83, p = 0.16). The three traits did not change independently (Figure S1); there was a significant positive correlation between the change in drug resistance and the change in growth rate (drug resistance decreased as growth rate decreased; Pearson correlation test, t_33_ = −3.68, p = 0.0008) and a significant negative correlation between drug tolerance and growth rate (as tolerance increased growth rate decreased (t_33_ = −2.39, p = 0.022; the correlation between change in drug tolerance and change in drug resistance was not significant, t_33_ = −0.30, p = 0.77). Thus, although the drug dosing regimen influenced the magnitude of some evolved phenotypic changes, across the evolutionary drug environments and transfer durations, the overarching pattern was that drug tolerance increased while growth rate (and, to a lesser extent, drug resistance) decreased in a correlated manner.

**Figure 2.**
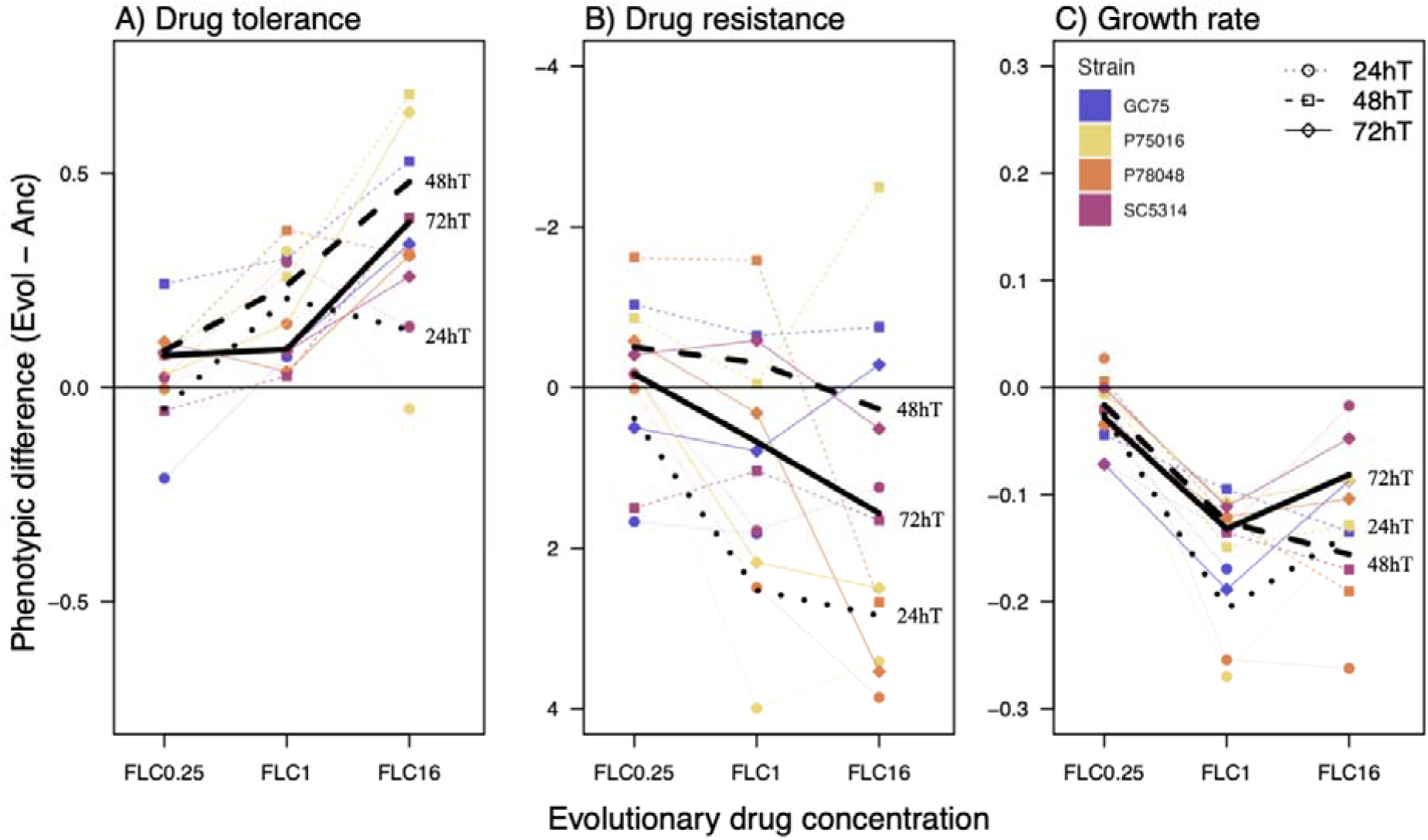
Phenotypic difference in evolved replicates compared to ancestral replicates. **A)** Drug tolerance, measured as FoG80 in a disk assay, B) drug susceptibility measured as RAD80 in a disk assay, C) growth rate measured in the evolutionary drug environment. Note that the drug reistance axis is flipped, to reflect that positive numbers indicate a larger zone of inhibition, i.e., a decrease in drug resistance Each point represents the mean of surviving evolved replicates from each strain background. The different point shapes indicate replicates with different transfer durations. The lines are for visualization purposes; coloured lines for each strain background with the thick black lines showing the overall mean for each drug concentration across all strains for each transfer regimen.

Increases in tolerance were pervasive across the drug-dosing regimens. The exception was 24 h transfers at FLC0.25, where three strain backgrounds showed no change, and evolved replicates from GC75 had decreased tolerance (Figure S2). The only other exceptions were the SC5314 populations evolved with 48 h transfers in FLC0.25, where the ancestral replicates had a modest but significantly higher tolerance, and FLC1, where there was no difference. Changes in resistance at the population level were less consistent. The majority of populations either exhibited no change or a reduction in resistance (Figure S3). At the population level, small yet significant increases between evolved and ancestral replicates were observed in GC75, P75016, and P78048 evolved with 48 h transfers in FLC0.25, and P78048 evolved with 48 h transfers in FLC1. A small number of individual replicates evolved to FLC16 from GC75 (with both 48 h and 72 h transfers), and P75016 (with 48 h transfers) evolved high resistance (Figure S3). Nearly all isolates evolved at all drug levels at all transfer durations decreased in growth rate (Figure S4).

To examine cross-tolerance and cross-resistance patterns in additional azole drugs, we focused on GC75 and SC5314 lines evolved through 72 h transfers to FLC0.25 and FLC16. The results were very similar to what we observed in fluconazole. Replicates from both strain backgrounds evolved in FLC16 significantly increased tolerance and decreased resistance relative to ancestral replicates in all four tested drugs, with only a few exceptions (Figure 3, full statistical results reported in Figures S3A & S4A). By contrast, replicates evolved in FLC0.25 from both strain backgrounds generally had the same phenotype as ancestral replicates (Figure 3, statistical results in Figures S3A & S4A).

**Figure 3.**
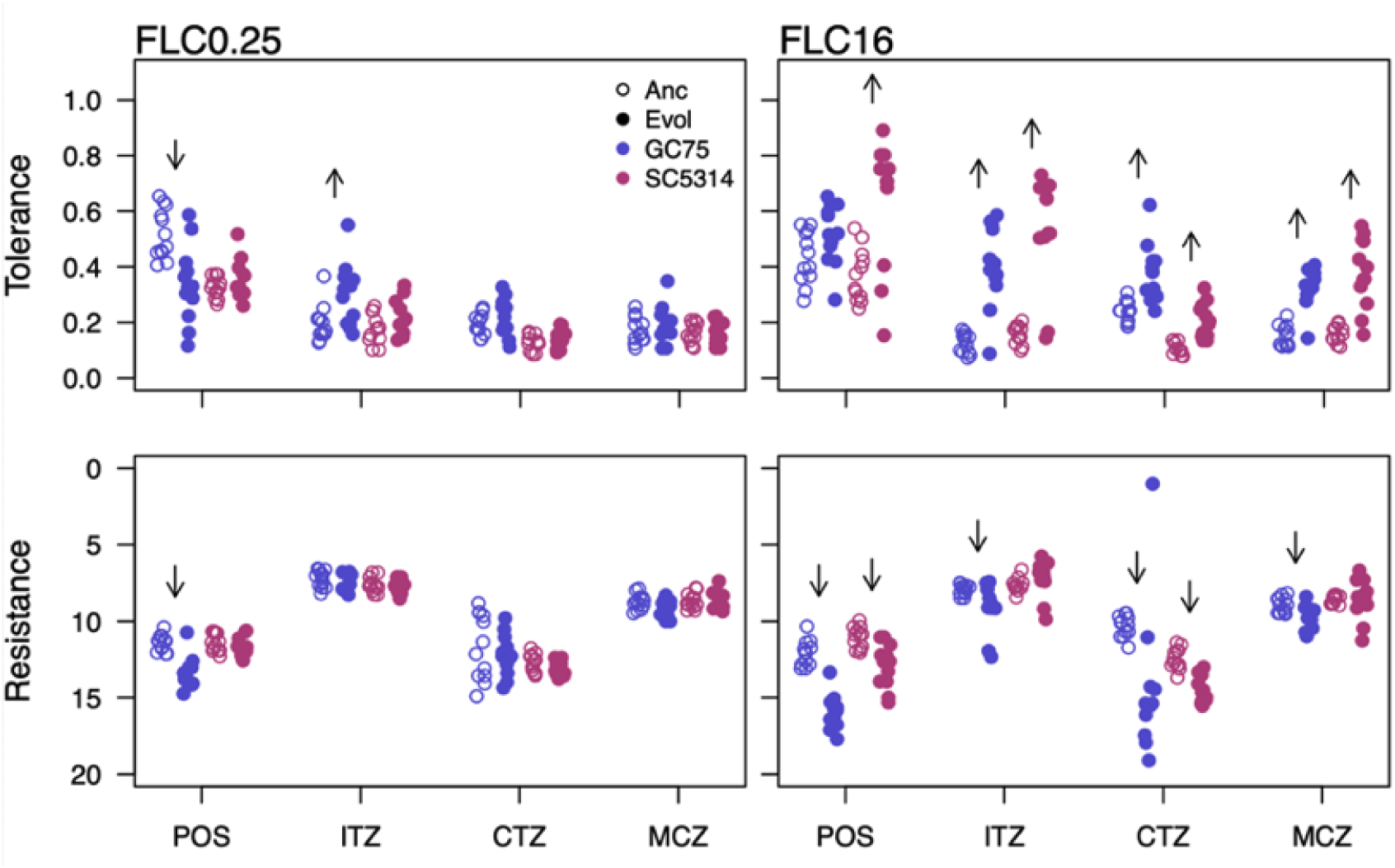
Drug response phenotypes measured in additional azole drugs. Tolerance (top) and resistance (bottom) of replicates from GC75 and SC315 evolved to FLC0.25 (left panels) and FLC16 (right panels) with 72 h transfers were examined for four drugs: posaconazole (POS), itraconazole (ITR), clotrimazole (CTZ), and miconazole (MCZ). Disk assay were conducted using the same methods as used for fluconazole. Each dot represents a single replicate.

Genome size increases have repeatedly been found in replicate lines evolved to fluconazole *in vitro*. In this short-term evolution experiment, we found that drug concentration, but not transfer duration (nor their interaction) was significantly associated with evolved genome size (Figure 4; linear mixed effect model treating strain as a random effect: drug concentration, F_2, 362.7_ = 13.66, p < 0.0001; transfer time, F_2, 362.7_ = 2.60, p = 0.08; drug × transfer time, F_4, 362.7_ =0.62, p = 0.69). Fewer than 3% of replicates evolved in FLC0.25 were above diploid, compared with 49% in FLC1 and 40% in FLC16. To examine whether evolved genome size was a significant predictor of drug tolerance or resistance, we ran linear mixed-effect models (one for each drug response phenotype) with replicate genome size, transfer duration, and evolutionary drug concentration as predictor variables in models that included the three single factors as well as all possible interactions and strain as a random effect. Drug tolerance was significantly influenced by all three predictors, with no significant interactions (shown is only the reduced model without interactions: drug concentration, F_2, 339..5_ = 117.42, p < 0.0001; transfer time, F_2,339.6_ = 45.64, p = 0.006; genome size, F_1,340.8_ =6.01, p = 0.015; Figure 5A). By contrast, drug resistance was significantly influenced by a three-way interaction among the three predictor variables (F_4,339.5_ = 26.8, p < 0.0001; Figure 5B). Given that, the single factors are difficult to interpret. A visual inspection shows no clear association between genome size and resistance; the few isolates that increased in resistance all remained diploid. Looking at the simplest possible model that ignores potentially confounding variables paints the same picture: there is a significant positive correlation between evolved genome size and tolerance (t_346_ = 3.66, p = 0.0003) but not between genome size and resistance (t_346_ = 0.68, p = 0.50).

**Figure 4.**
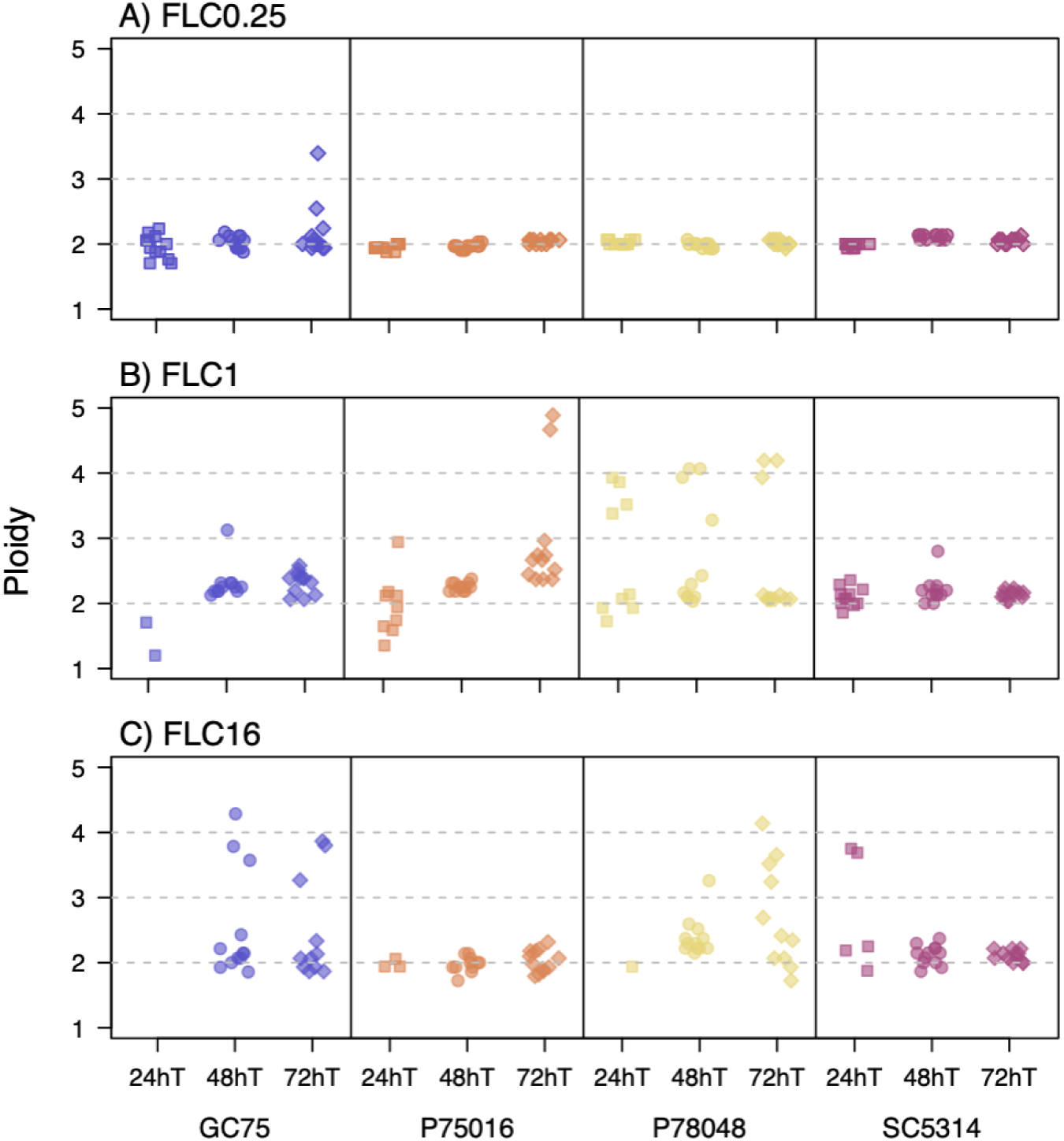
Genome size was commonly above diploid in evolved isolates. Ploidy was calculated by dividing evolved fluorescence intensity of evolved replicates by the median fluorescence intensity of ancestral replicates from the same strain background, evolved drug concentration and transfer duration (ancestral and evolved replicates were always quantified in the same experimental batch).

**Figure 5.**
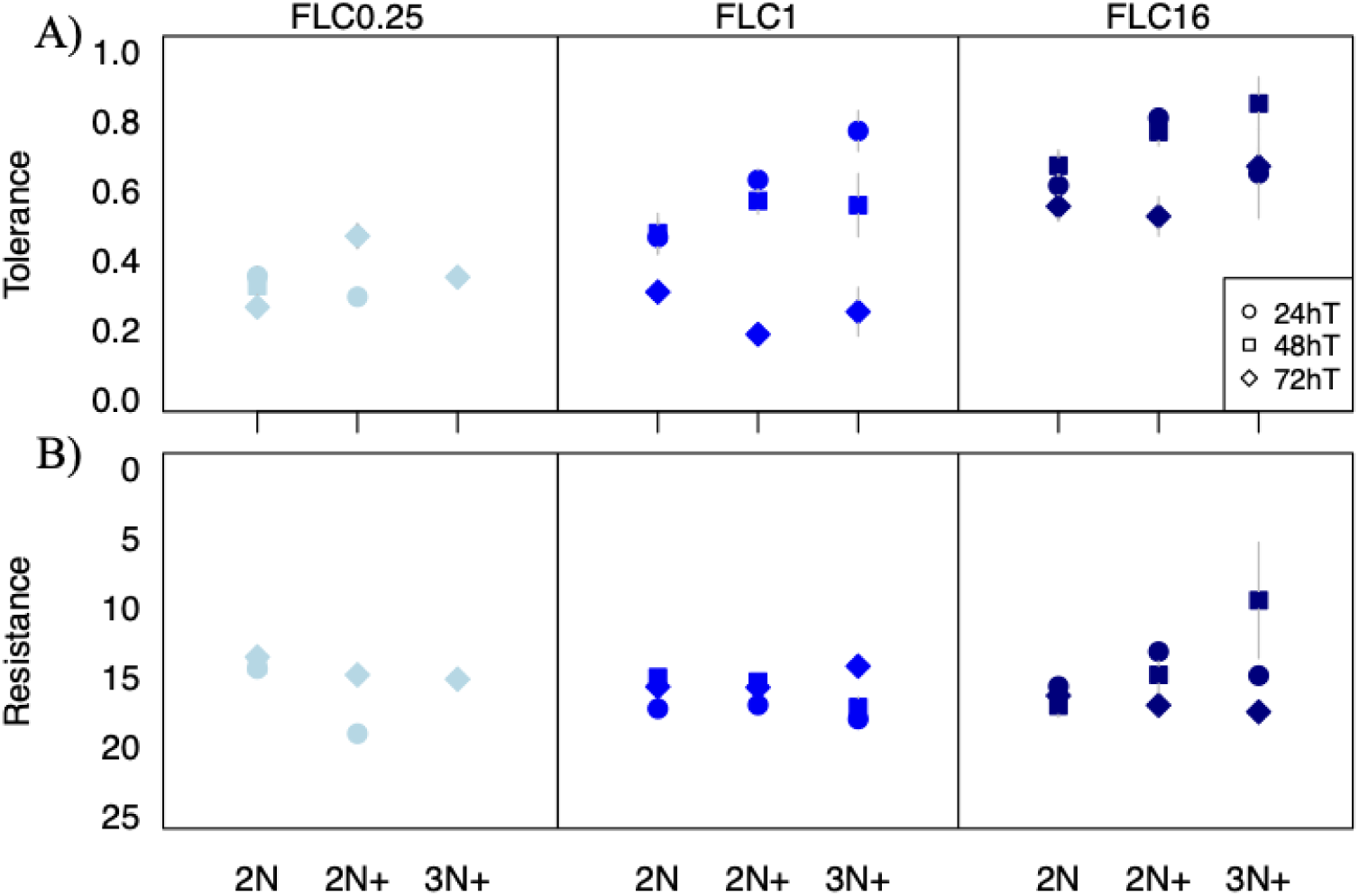
Evolved genome size was significantly associated with evolved tolerance but not resistance. Quantitative ploidy bins were determined from ancestral replicate data; as all ancestral replicates are diploid, the cutoff for 2N+ was set based on the value (2.2) that corresponded to fewer than 1 % of ancestral replicates being classified as larger than diploid. The 3N+ bin was correspondingly set at 3.2. Error bars represent the standard error of all isolates in each group, irrespective of strain background.

## DISCUSSION

The acquisition of antimicrobial resistance is an inherently evolutionary process. Drug-susceptible populations of microbes acquire mutations. In the presence of drug, the beneficial mutations are rapidly selected, and the proportion of cells carrying those mutations increases in frequency. In the presence of fungistatic drugs, such as the commonly prescribed antifungal fluconazole, selection could theoretically act not only on beneficial mutations that increase drug resistance, but also on mutations that increase drug tolerance, the proportion of the population that is able to grow slowly at drug levels above the inhibitory concentration (MIC) (Berman & Krysan, 2020; Gerstein et al., 2016; Gerstein & Sethi, 2022; Rosenberg et al., 2018). *In vitro* evolution experiments have been used to examine how different factors such as strain background and evolutionary drug concentration influence the propensity for evolved replicates to acquire increases in resistance and/or tolerance. Here, we conducted a fully factorial experiment with four *C. albicans* strain backgrounds to examine the combined influences of drug concentration (sub-MIC_50_, at/near MIC_50_, and supra-MIC_50_) and the duration of time between transfer into fresh drug medium (24 h, 48 h, 72 h) on short-term adaptation to fluconazole. We collectively refer to these factors as the drug-dosing regimen. Our overarching results across the nine tested regimens indicated that increases in fluconazole tolerance (and cross-tolerance to other azoles) were pervasive in evolved replicates from all strain backgrounds, alongside correlated increases in genome size and decreases in growth rate. By contrast, increases in resistance were rare, indicating that, in this batch-culture experimental framework, selection on mutations that increase tolerance is the most common evolutionary trajectory.

Our *a priori* expectation was that replicates evolved with a 24 h transfer duration, particularly at higher drug levels, would promote selection for resistance over tolerance. Drug resistance is observed (and measured) at 24 h, while drug tolerance requires growth quantification (in the same assays as used for resistance) at or after 48 h (Berman & Krysan, 2020; Gerstein et al., 2016; Rosenberg et al., 2018). We reasoned that there should be no scope for tolerance selection in the 24 h transfer experiments, which would push populations towards selection for increased resistance. However, this was not borne out. Replicates transferred after 24 h at all three drug levels showed relatively small increases in tolerance compared with the other transfer times, yet they also showed the largest average decrease in resistance compared with replicates evolved with a longer transfer duration. Resistance and tolerance seem to be governed by independent genetic processes. Several laboratory evolution experiments have documented increases in drug tolerance during experimental evolution to azoles, alongside a corresponding increase in genome size, and decrease (or no change) in drug resistance (Gerstein & Berman, 2020; Guo et al., 2024; Kukurudz et al., 2022; Sun Liu-liu et al., 2023; Todd et al., 2023; Yang et al., 2023).

Chromosome R aneuploidy appears to be the most common genetic route to increased tolerance in batch culture evolution experiments (Guo et al., 2024; Kukurudz et al., 2022; Sun Liu-liu et al., 2023; Todd et al., 2023; Yang et al., 2023). A recent study using high-throughput CRISPR activation and CRISPR inhibition across all R chromosome genes showed that overexpressing *SEC4* and *YPT31* (located ∼1200 kb apart on the left arm) is both necessary and sufficient to confer tolerance increases (Gervais et al., 2026). Chromosomal aneuploidy occurs at a higher mutation rate than the per-base-pair mutation rate (Ene et al., 2018; Zhu et al., 2014); if a beneficial aneuploidy is available with a sufficiently high effect size, it will be the most common outcome of the partially stochastic process of natural selection (Gerstein & Berman, 2015). Common laboratory environments do not well mimic the human body, however. Since batch culture evolution experiments are predicated on the transfer of a specific ratio of cells (typically 1:00 or 1:000), in *C. albicans* they are almost all done at 30 °C rather than 37 °C and in rich medium, to deliberately keep fungal cells in the yeast rather than hyphal phase. Whether chrR aneuploidy also appears readily in populations in the human body, and thus, whether this is a relevant result to understanding evolution “in the wild”, is not known. It may be that the tolerance benefits conveyed by chrR aneuploidy in the body are tempered by the deleterious effects of increasing the copy number of the 992 other genes on that chromosome. Approximately 9.5% of previously whole-genome sequenced *C. albicans* isolates are aneuploid (Adamu-Bukari, *et al*., 2026), though a recent review found that the prevalence of copy number variants was much higher in *in vitro* experiments than in natural isolates (Jay et al., 2025). Many different mutations across the genome that provide increases in drug resistance (rather than tolerance) have been identified from clinical isolates (Bédard et al., 2025). It remains an open question how often tolerance is involved in clinically recalcitrant infections and the role of aneuploidy in nature.

The other overwhelming phenotypic change we observed across dosing regimens was a decrease in growth rate, which correlated with increases in tolerance. This was most pronounced in the higher drug levels, with no clear pattern between different transfer durations. Returning again to the definition of drug tolerance, this makes sense—a hallmark of the subpopulation of tolerant cells is slow growth in supra-MIC drug concentrations (Berman & Krysan, 2020). The plate reader growth assay we used captures only the population growth average. Single-celled approaches are required to capture and quantify growth rate from the two populations of cells (i.e. the tolerant and non-tolerant populations), such as seen in the ScanLag method that differentiates tolerant and non-tolerant subpopulations through the timing of colony appearance (Gautier et al., 2025; Levin-Reisman et al., 2010; Rosenberg et al., 2018).

The most obvious influence of the drug dosing regimen was the strong association with replicate extinction, which was observed only in the 24 h transfer experiments. The extinction rate was strongly positively correlated with evolutionary drug concentration and associated with strain background. GC75 strain background replicates were the most likely to go extinct across all drug concentrations, with SC5314 replicates the least likely to go extinct (and P75017 and P78048 intermediate). Two previous evolution experiments we conducted in different drugs (posaconazole and boric acid) also found that strain background was significantly associated with extinction. Interestingly, however, different drugs yielded different patterns: in posaconazole, replicates initiated from P75016 were the most likely to go extinct, followed by GC75, then SC5314 and P78048; in boric acid, in contrast to the azoles, SC5314 replicates went extinct the fastest, while P75016 and P78048 replicates went extinct the least. Presumably, there are strain-background associated differences in the spectrum of available mutations that can rescue the population from extinction. A different experimental framework with sufficient statistical power, purposefully designed to explore the genomic factors that influence extinction probability, will be required to identify genomic correlates. Future experiments focused on extinction should be also be done purposefully in environments that are known to influence drug tolerance, such as temperature, growth medium, and pH (Agrawal et al., 2007; Coenye et al., 2008; Delarze et al., 2020; Marr et al., 1999; Yang et al., 2023). If baseline levels of drug tolerance are reduced through abiotic factors, the rate of extinction should increase.

Combined, our work highlights that increases in drug tolerance, rather than drug resistance, repeatedly evolved in replicate lines of C. albicans in short-term experiments, evolved to different fluconazole levels and transfer durations. We also demonstrate that with evolution to fluconazole, growth rate is a poor proxy for fitness, as decreases in growth rate were observed across all regimens and significantly correlated with increases in drug tolerance. Furthermore, we found that replicate extinction was very common with 24 h transfers, with the rate influenced by drug concentration and strain background. We hope our results motivate future work to report replicate extinction more comprehensively and to design purposeful experiments that explore abiotic and genomic factors and their influence on extinction directly.

