## Supplemental Figures 1-4 for "The influence of drug dosing regimen on extinction and adaptation during experimental evolution in *Candida albicans*"

**SUPPPLEMENTAL FIGURES**

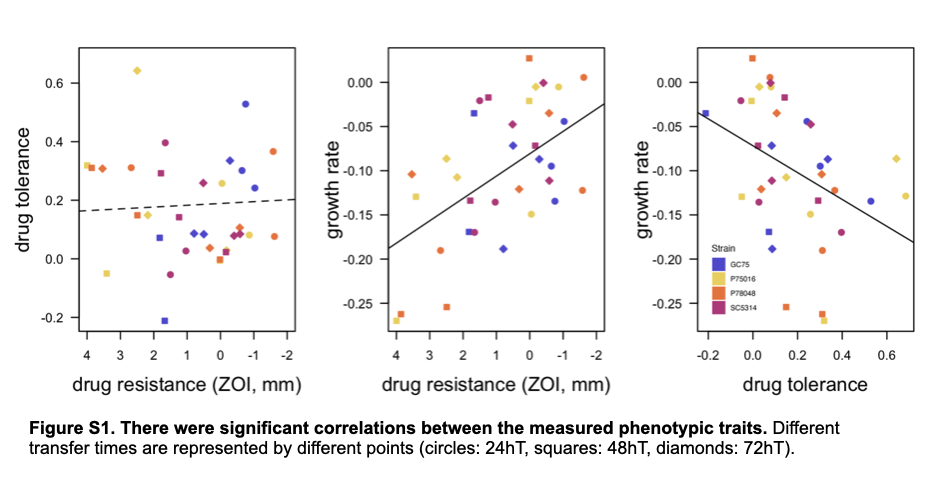

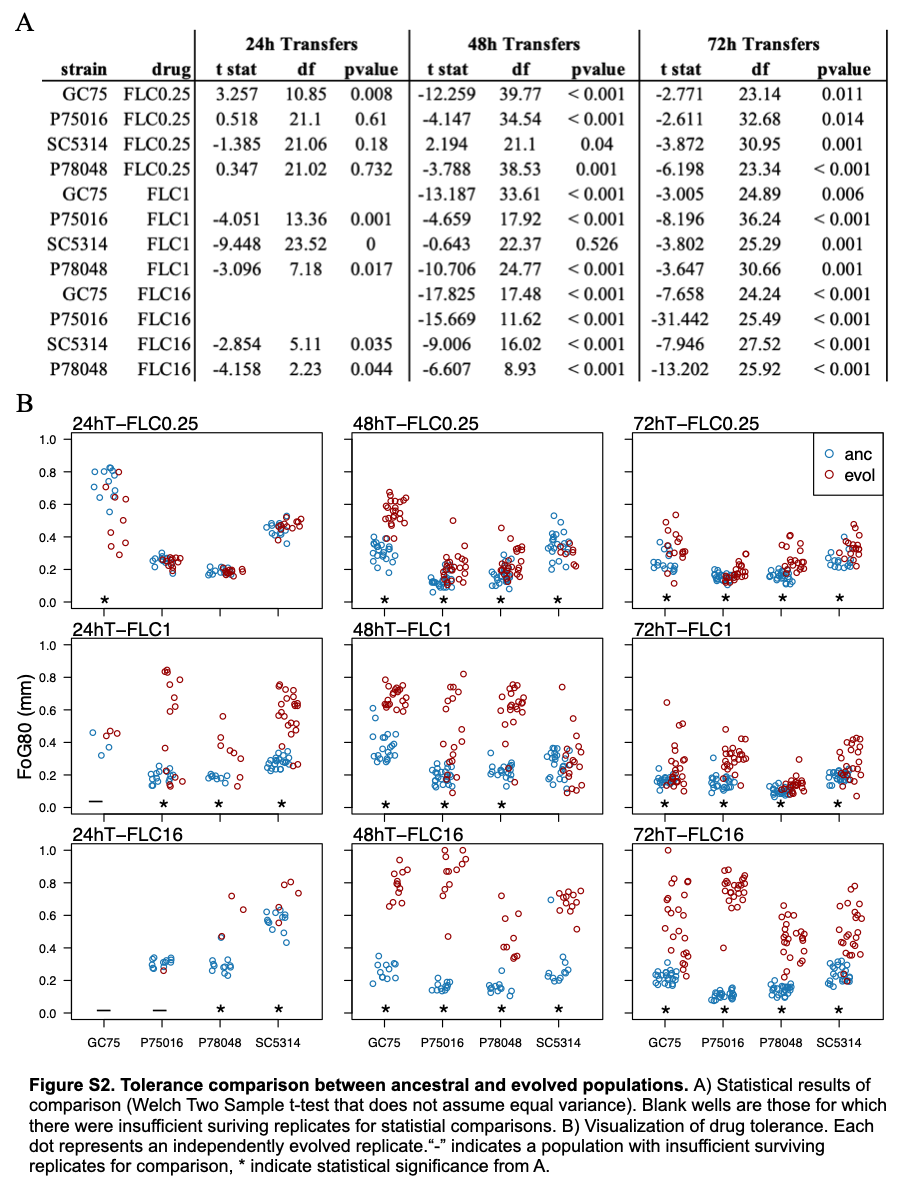

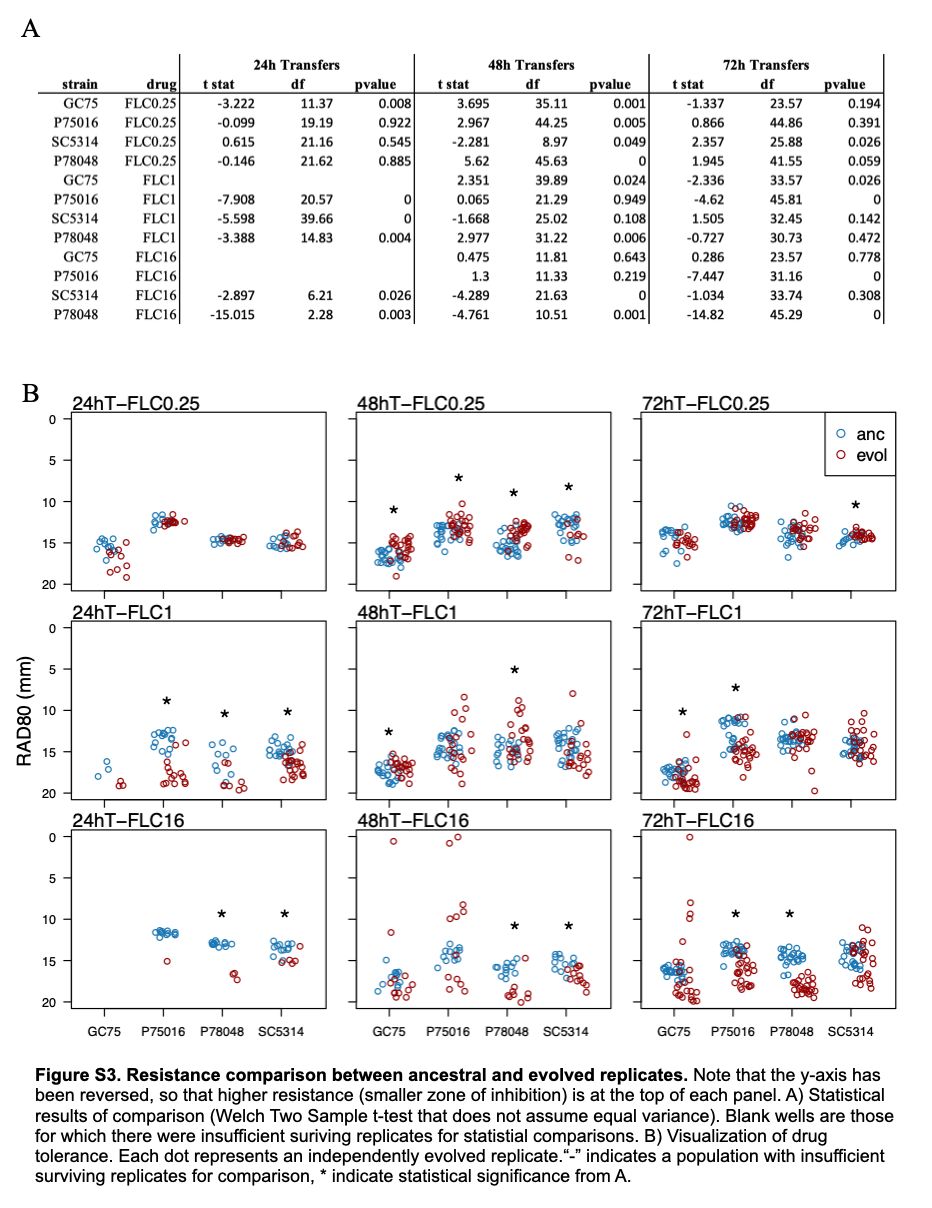

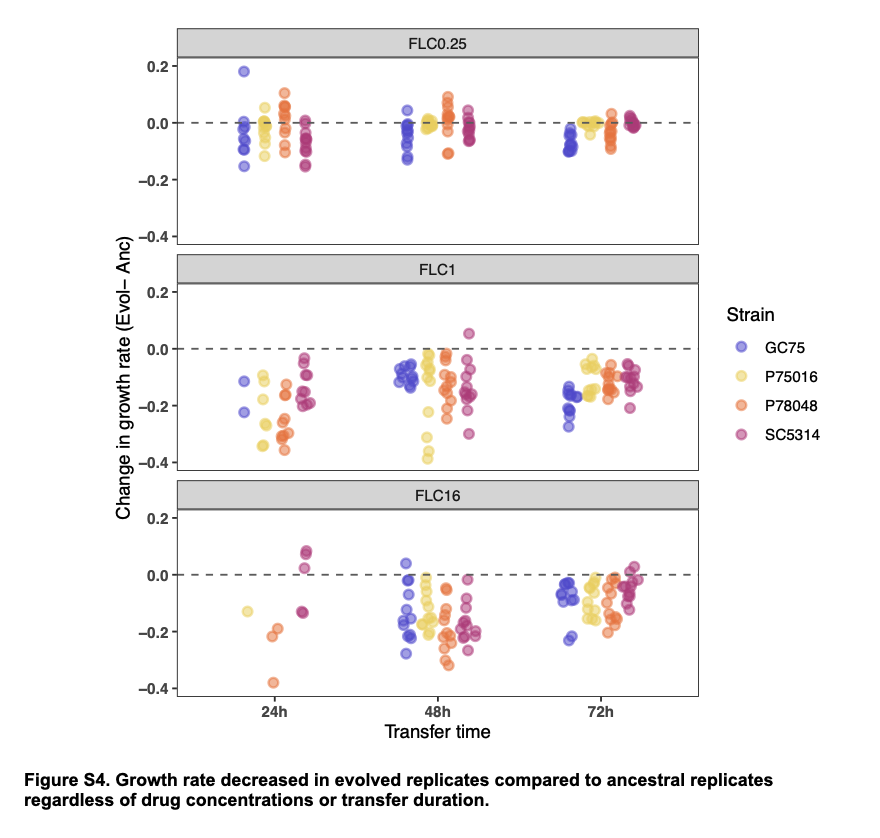
